# Biocompatible PVDF Nanofibers with Embedded Magnetite Nanodiscs Enable Wireless Magnetoelectric Neuromodulation

**DOI:** 10.1101/2025.06.16.660052

**Authors:** Lorenzo Signorelli, Anouk Wolters, Vicente Duran Toro, Elif Koҫar, Franziska Wasner, Nadine I. Goldenstein, Jonas Englhard, Mahdieh Shojaei Baghini, Hadi Heidari, Julien Bachmann, Sarah Hescham, Danijela Gregurec

## Abstract

Wireless neuromodulation technologies aim to eliminate the need for invasive hardware and enhance tissue compatibility. Magnetoelectric (ME) materials enable magnetic field-induced electrical stimulation, offering a minimally invasive neural activation. However, conventional ME systems use rigid ceramic components with limited biocompatibility. Here, we report a flexible, predominantly organic ME platform composed of polyvinylidene fluoride (PVDF) nanofibers embedded with anisotropic magnetite nanodiscs (MNDs). These MNDs were selected for their unique ability to exert magnetic torque due to vortex magnetization, and their intrinsic magnetostrictive behaviour. The resulting ME fibers preserve the piezoelectric β-phase of PVDF and exhibit magnetoelectric voltage coefficient of 1.26 Vcm⁻¹Oe⁻¹. We compare two magnetic activation strategies; torque-based and high-frequency magnetostriction, finding that magnetostriction more effectively triggers neuronal responses. In vitro calcium imaging reveals robust activation in primary cortical neurons cultured on ME fibers. Biocompatibility post-stimulation was confirmed on ex vivo human brain tissue, with no increased cell death. Implanted into the premotor cortex of freely moving mice, the fibers enabled wireless modulation of motor behaviour under an alternating magnetic field. This work presents the first demonstration of wireless magnetoelectric neuromodulation using soft, biocompatible fiber composites, paving the way for future bioelectronic interfaces free from rigid components and tethered systems.

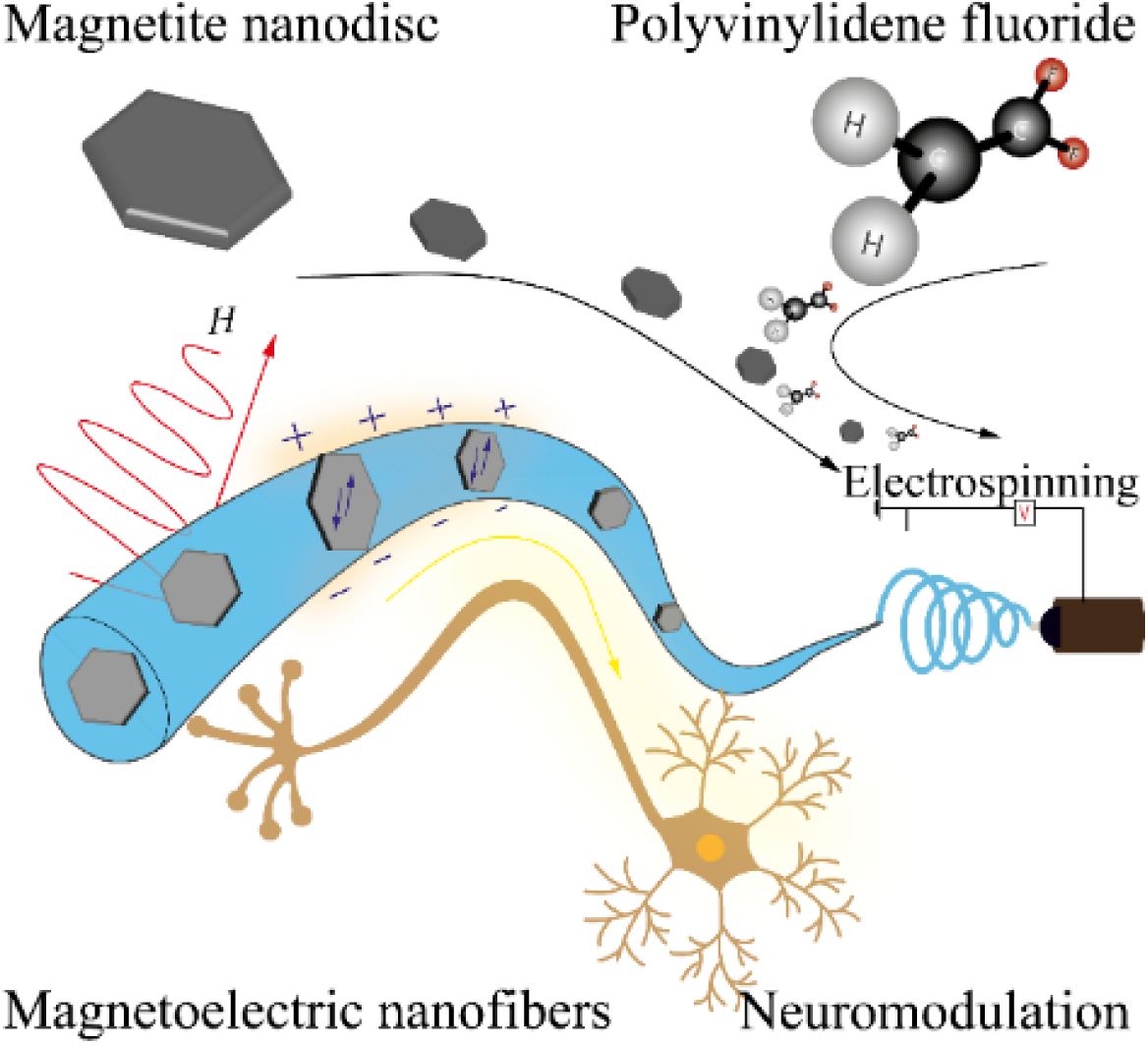

## 1. Introduction

Magnetoelectric materials which convert magnetic stimuli into electrical output via mechanical coupling, are gaining momentum in bioelectronics and neuromodulation applications due to their ability to deliver remote, localized stimulation without physical wiring[1, 2]. These materials typically exploit the coupling between magnetostrictive and piezoelectric phases, enabling magnetic field-induced mechanical strain to produce an electric response[2]. To date, the most common ME nanostructures employ core-shell geometries combining cobalt ferrite (CoFe_2_O_4_) and barium titanate (BaTiO_3_)[3]. These systems achieve high ME coefficients and have demonstrated efficacy in modulating neuronal activity both in vitro and *in vivo*[1]. However, the use of cobalt and barium raises concerns about biocompatibility, especially for chronic or implantable applications. Studies report potential neurotoxicity and systemic effects from long-term exposure to degradation products of these materials[4–7] underscoring the need for safer alternatives. Polyvinylidene fluoride (PVDF) has emerged as a promising candidate for next-generation soft ME systems. This semi-crystalline polymer is inherently piezoelectric in its β-phase and offers excellent flexibility, chemical inertness, and biocompatibility[3, 8]. Electrospinning of PVDF promotes β-phase formation through both mechanical stretching and electric field-induced chain alignment, significantly enhancing its electromechanical properties[9, 10]. Additionally, traditional neurostimulation electrodes such as Utah Arrays are rigid and bulky, with a Young’s modulus in the GPa range, which can damage surrounding tissue and lack stimulation precision due to the widespread electric fields[9, 11, 12]. PVDF, though also stiff as bulk material (2.5–3.5 GPa), becomes highly flexible at the nanoscale; electrospun PVDF nanofibers exhibit a much lower modulus (~86 MPa), closer to brain tissue (~1 kPa). Given its biocompatibility and piezoelectric β-phase properties[13] PVDF nanofibers enriched with anisotropic Fe_3_O_4_ magnetite nanodiscs[14] offer a promising, minimally invasive alternative for targeted, magnetoelectrically driven neurostimulation.

In this work, we introduce a predominantly organic ME nanofiber platform composed of electrospun PVDF anisotropic magnetite nanodiscs (MNDs). These MNDs, synthesized via a template-assisted reduction method[14], exhibit a vortex magnetization ground state and are capable of undergoing controlled magnetization reorientation under applied magnetic fields. Their disc-like shape imparts high magnetic anisotropy, enabling two forms of actuation: (i) torque generation at low (~5 Hz) frequencies via vortex-to-in-plane transitions, and (ii) magnetostrictive deformation at higher (~150 Hz) frequencies combined with the static field[14, 15] (**Figure 1**). Previous work has shown that shape anisotropy significantly influences the magneto–mechano–electrical response of magnetic piezoelectric composites. For instance, Kim et al. demonstrated that nanorod-decorated P(VDF-TrFE) fibers produced significantly higher electrical output than spherical nanoparticles under alternating magnetic fields, due to enhanced magnetic torque[16]. However, their system focused on energy harvesting and relied on surface decoration rather than full embedding. By contrast, our approach embeds anisotropic magnetite nanodiscs directly within the PVDF matrix, which promotes more efficient strain transfer, improved mechanical stability, and enhanced biocompatibility. Although bulk magnetite exhibits only moderate magnetostriction, its anisotropic nanoscale form significantly amplifies strain transfer due to shape-enhanced magnetic alignment as recently reported by Kim et al[17], achieving high ME effect due to the anisotropy of the ferrites and the double core-shell structure based on magnetite and cobalt ferrite (Fe_3_O_4_–CoFe_2_O_4_). We hypothesized that this anisotropy-enhanced magnetostrictive effect of anisotropic MNDs, when coupled with the piezoelectric response of PVDF, could produce magnetoelectric outputs sufficient for neuronal stimulation.

**Figure 1.**
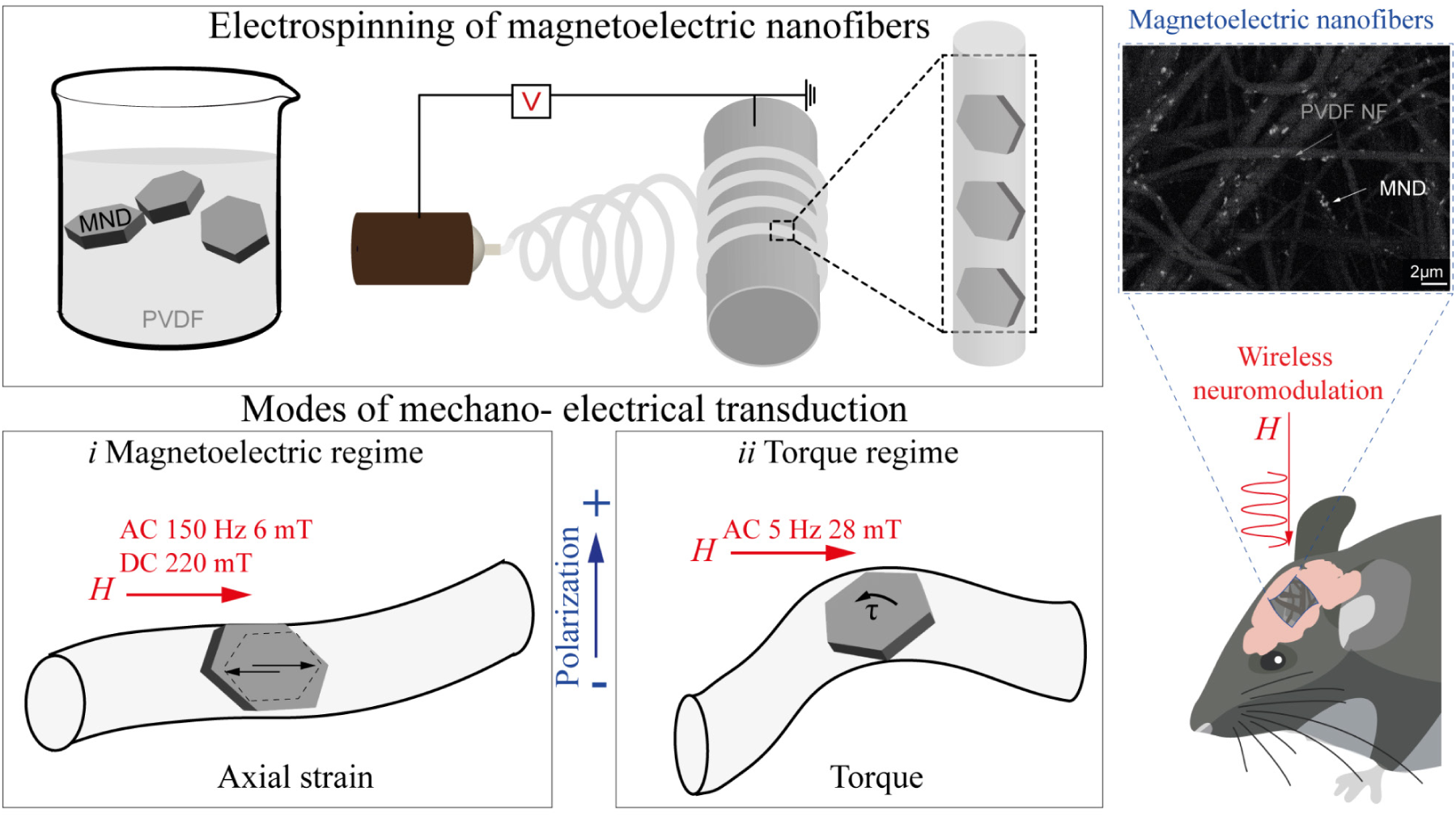
Magnetoelectric nanofibers (MEFs) are prepared by electrospinning embedding the magnetite nanodiscs (MNDs) within the PVDF fiber. Mechano-electrical transduction can occur through two modes: *i* Under a combined static (DC 220 mT) and alternating (AC 6 mT, 150 Hz) magnetic field, MNDs undergo domain oscillations that generate axial strain in the surrounding piezoelectric PVDF matrix. This mechanical deformation induces electric potential via the piezoelectric effect. *ii* At low-frequency magnetic fields (AC 26 mT, 5 Hz), the vortex-to-in-plane magnetization transition in the MNDs produces internal torque. This torque twists the PVDF nanofiber, leading to lateral deformation and electric polarization. In both cases, the embedded configuration enables efficient mechanical coupling between the MNDs and the PVDF matrix, allowing wireless, strain-mediated voltage generation. Right panel shows a high contrast SEM micrograph of nanofibers embedding MNDs, along with a schematic illustrating the implantation of MEFs on the surface of the mouse brain for magnetic field-mediated wireless neuromodulation.

To evaluate this, we fabricated magnetoelectric fibers (MEFs) using electrospinning and characterized their morphology, crystallinity, magnetic properties, and ME coupling. We first compared two actuation modes: torque-based and magnetostrictive stimulation in primary cortical neurons using calcium imaging. The magnetostrictive mode elicited robust neuronal activation, which was significantly reduced by tetrodotoxin (TTX), confirming the involvement of voltage-gated ion channels. In contrast, torque-based stimulation showed no significant neuromodulatory effect. Biocompatibility was confirmed using *ex vivo* human brain tissue, and MEFs implanted in the premotor cortex of freely moving mice modulated motor behavior upon wireless magnetic stimulation via magnetostrictive approach.

This study presents the first demonstration of wireless magnetoelectric neuromodulation using a fully soft, biocompatible fiber composite. By leveraging shape anisotropy in magnetite nanodiscs, we unlock effective strain-mediated activation in a flexible polymeric matrix, providing a scalable and safe platform for minimally invasive bioelectronic interfaces.

## 2. Results and Discussion

### 2.1. Synthesis and Characterization of Magnetoelectric Nanofibers

Anisotropic magnetite nanodiscs (MNDs) were synthesized using our previously reported template-assisted reduction method[14], yielding uniform hexagonal structures approximately 230–240 nm in diameter (**Figure 2a**). These nanodiscs exhibit a vortex magnetization ground state and can undergo controlled vortex-to-in-plane transitions under low-frequency (<10 Hz) magnetic fields[14, 17]. The anisotropic geometry of the MNDs is also known to enhance magnetostrictive behavior compared to isotropic faceted nanoparticles, enabling efficient magnetomechanical coupling[17]. The MNDs were structurally confirmed via X-ray diffraction (XRD), which showed a pure magnetite phase (Figure S1). Vibrating sample magnetometry (VSM) revealed a saturation magnetization (Ms) of approximately 75 emu/g, closely matching the value for bulk magnetite[15].

**Figure 2.**
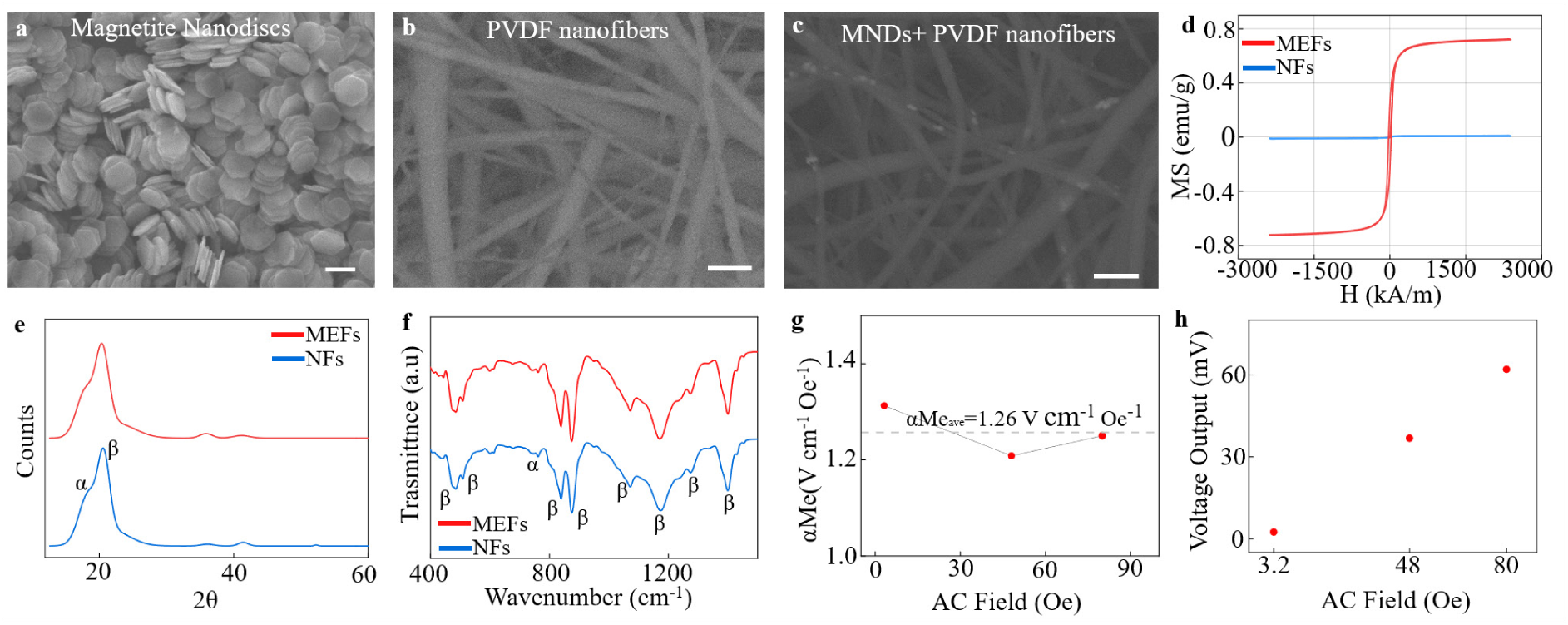
Synthesis and Characterization of magnetoelectric nanofibers: a) SEM micrograph of magnetic nanodiscs synthesized through solvothermal synthesis, then reduced through reflux of H_2_ gas. Scale bar: 200nm. b) SEM micrograph of pure PVDF nanofibers. c) Micrograph of nanodiscs embedded in PVDF nanofibers Scale bar: 2μm d) Magnetic hysteresis curves of PVDF and MEFs, showing preserved magnetic properties of MNDs after embedding. e) XRD spectra show the presence of both α and β phases in PVDF. f) FTIR spectra of MEF show characteristic peaks of the β phase at 838, 1072, 1170, and 1233 cm¹, g) Magnetoelectric coefficient of MEFs concerning the applied magnetic field shows an average αMe of 1.26 V cm¹Oe¹. h) Linear increase of the output voltage concerning the magnetic field at 120 Hz.

To fabricate magnetoelectric fibers (MEFs), the MNDs were incorporated into polyvinylidene fluoride (PVDF) nanofibers via electrospinning (Figure S2). A 0.1% (w/w) dispersion of dry MNDs was added to a 16% PVDF dispersion in DMF:acetone (1:1) and electrospun to form uniform, defect-free fibers (Figure S3). Scanning electron microscopy (SEM) confirmed that MND incorporation did not alter the fiber morphology or average diameter (Figure 2b–d and Figure S4). XRD analysis of the MEFs confirmed the presence of both α- and β-phases of PVDF. The β-phase, essential for piezoelectricity, was evidenced by the diffraction peak at 2θ = 21.88°, corresponding to the (200) reflection. Fourier-transform infrared spectroscopy (FTIR) corroborated these findings, showing characteristic β-phase absorption peaks at 474[18], 506[19], 840[20], 877[21], 1071[22], 1275[23], and 1401[10, 24] cm⁻¹, while a distinct peak identifies the α-phase at 762[25] cm⁻¹ (Figure 2e–f). The presence of the β-phase is primarily attributed to mechanical stretching during electrospinning, as well as the alignment of PVDF chains under the applied electric field[10]. These results align with previous reports showing enhanced β-phase formation in electrospun PVDF nanofibers[26]. Magnetization hysteresis measurements confirmed that MNDs retained their magnetic properties within the polymer matrix, showing characteristic ferromagnetic behavior in MEFs but not in pure PVDF fibers (Figure 2g). This suggests that while Brownian motion is suppressed post-embedding, the MNDs still respond via Néel relaxation. To evaluate the magnetoelectric performance, we quantified the output voltage of MEFs under varying magnetic field strengths (Figure S5). Signals from pure PVDF NFs, likely arising from environmental electromagnetic noise or mechanical perturbation, were subtracted as baseline. The corrected voltage output was normalized by fiber thickness (~500 nm) (Figure S6) to obtain the ME voltage coefficient (αME), which averaged 1.26 V·cm⁻¹·Oe⁻¹ (Figure 3h). A linear increase in voltage with applied field was observed, consistent with piezoelectric strain transduction.

**Figure 3:**
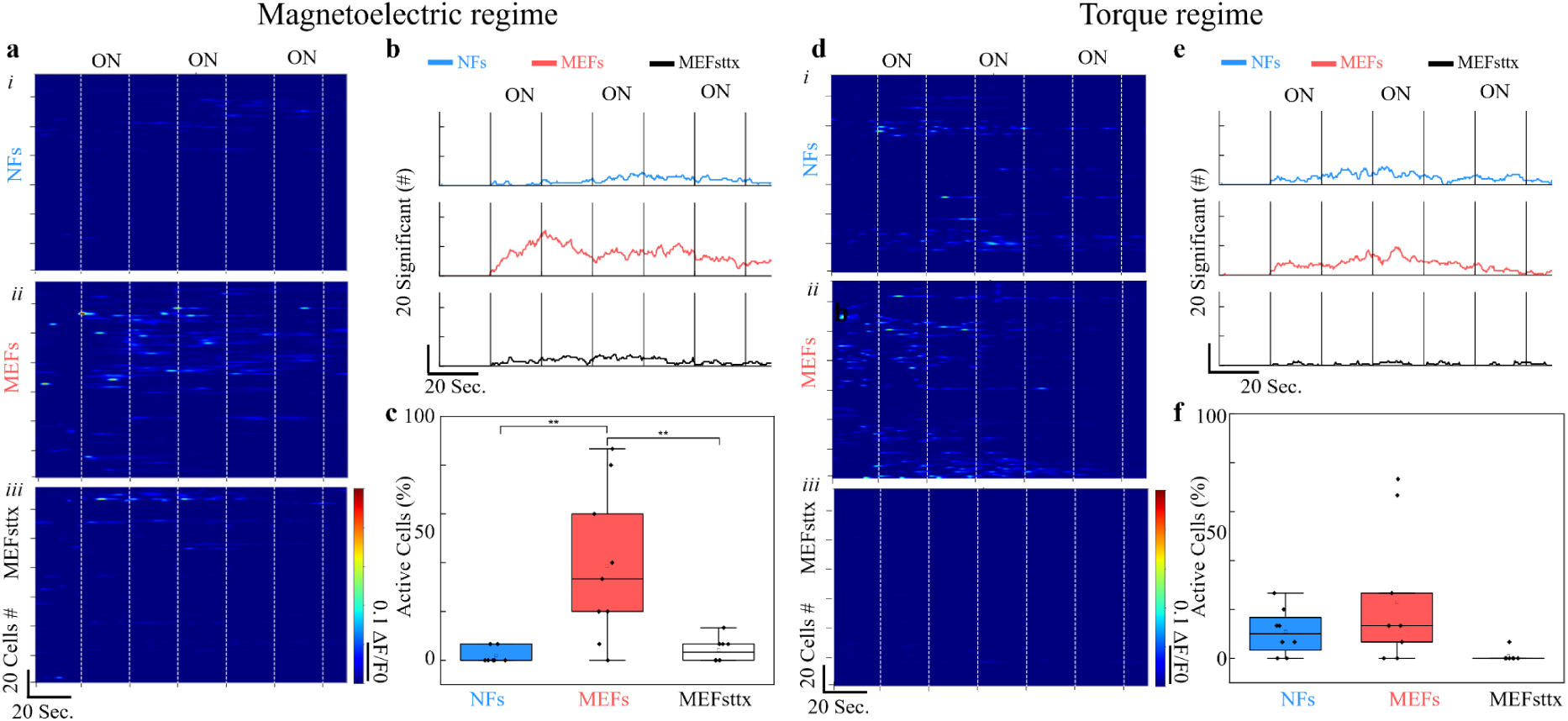
Primary cortical neurons show a significant increase in activation when stimulated via magnetoelectric actuation, while they do not exhibit significant differences when the fibers are stimulated through torque; a) heat map under magnetelectric conditions for NFs (i), MEFs (ii), and MEFsttx (iii); b) cells showing significantly different activity compared to baseline; c) percentage of activated cells (p < 0.01); d) heat map under torque activation for NFs (i), MEFs (ii), and MEFsttx (iii); e) cells showing significantly different activity compared to baseline; f) percentage of activated cells

These results demonstrate that our fabrication approach successfully preserves the key functional properties of both components within the composite: the magnetic responsiveness of the anisotropic MNDs and the piezoelectric β-phase crystallinity of PVDF. The nanodisc geometry and uniform embedding ensure that magnetic stimuli are mechanically coupled to the surrounding polymer, enabling magnetostrictive strain transfer. Importantly, the electrospinning process not only aligns the PVDF chains to favor β-phase formation but also accommodates MNDs without disrupting fiber morphology or mechanical integrity.

The observed magnetoelectric voltage coefficient (α ME ≈ 1.26 Vcm⁻¹Oe⁻¹) is comparable to or exceeds that of many ceramic- or hybrid-based ME systems[2, 3], despite being entirely polymer-based. This is particularly significant given the biocompatibility, flexibility, and minimal toxicity of the materials used. Unlike brittle ceramic composites, these fibers retain mechanical robustness and processability at sub-micron scales, making them more suitable for implantable applications. In summary, the developed MEFs combine nanoscale magnetic anisotropy with efficient electromechanical conversion in a soft, biocompatible format bridging the gap between high-performance ME behavior and biomedical applicability.

### 2.2. *In Vitro* Testing of Magnetoelectric Activation Modalities and Neural Tissue Compatibility

To evaluate the neuromodulatory potential of MEFs, we conducted *in vitro* calcium imaging in primary cortical neurons and assessed material biocompatibility using *ex vivo* human brain slices. Together, these experiments aimed to validate both the functional efficacy and safety of the developed fibers under conditions relevant to neural stimulation. Primary cortical neurons isolated from neonatal rats were cultured for 10 days on either MEFs or pure PVDF NFs. On the day of imaging, neurons were incubated with Fluo-4 and Dooku1 (a PIEZO1 inhibitor) to reduce mechanical stimulation artifacts. Cells were then subjected to magnetic field stimulation using two modes: (i) magnetostrictive stimulation (220 mT DC + 6 mT AC @ 150 Hz), and (ii) torque-based stimulation (26 mT AC @ 5 Hz) (Figure S7).

Under magnetostrictive conditions, neurons cultured on MEFs showed robust calcium transients in response to magnetic pulses (Figure 3a(ii)). This effect was significantly attenuated in samples pre-treated with tetrodotoxin (TTX), confirming the involvement of voltage-gated ion channels (Figure 3a(iii)). In contrast, neurons on NFs exhibited no activity (Figure 3a(i)). Quantitative analysis showed significantly elevated calcium event frequency and the proportion of active neurons in MEF-stimulated cultures compared to controls (Figure 3b–c). Under torque-based stimulation, no significant differences in activity were observed across conditions (Figure 3d–f), indicating that torque alone was insufficient to elicit piezoelectricity and neuronal activation under the tested parameters.

To confirm the biocompatibility of MEFs and rule out the toxic effects of magnetic stimulation, we used human cortical brain slices obtained from resective epilepsy surgery. Tissue was incubated for 48 h after exposure to the same stimulation protocol (3 min, 120 mT DC + 6 mT AC at 150 Hz), with or without the presence of NFs or MEFs. Cell viability was assessed using propidium iodide staining, a marker of cell death. Quantitative fluorescence analysis revealed no significant increase in dead cells across any of the conditions or timepoints (**Figure 4a–d**). The safety of the MEF fibres was also evaluated on HEK cells over 6 days using the IncuCyte® S3 imager (Sartorius, USA) with the basic analyser module (Figure S8), where none of the components adversely affected cell health or viability during the observation period.

**Figure 4.**
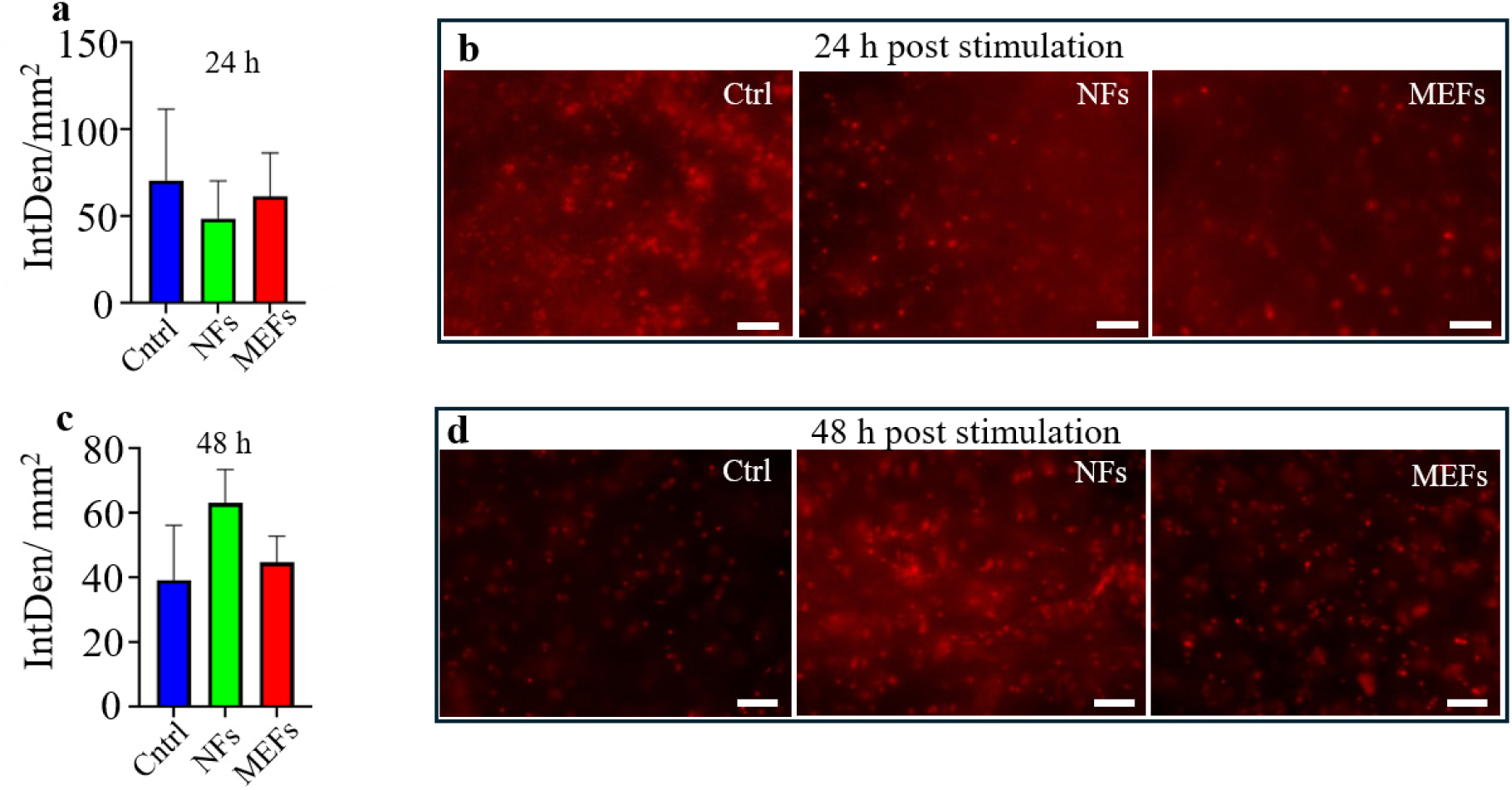
Biocompatibility assessment of PVDF nanofibers (NFs) and magnetoelectric fibers (MEFs) using resected human brain slices. a) Quantification of cell death via integrated propidium iodide fluorescence intensity following 24 h incubation after magnetic stimulation (120 mT DC + 6 mT AC, 150 Hz, 3 min). b) Representative fluorescence images of brain slices from control, NF-treated, and MEF-treated groups after 24 h. c) Integrated fluorescence intensity following 48 h incubation. d) Representative images of brain slices from the same groups after 48 h. No significant differences in cell death were observed between conditions at either time point.

Taken together, these findings demonstrate that among the two tested magnetoelectric stimulation strategies, magnetostrictive activation outperforms torque-based actuation in eliciting neuronal responses. The likely explanation is that at the tested frequencies and field strengths, magnetostriction results in more consistent and directional strain transfer to the PVDF matrix, leading to stronger and more uniform piezoelectric output. In contrast, torque-based deformation may be insufficient in magnitude or poorly transmitted through the fiber structure rendering it unreliable for activating voltage-gated ion channels. Based on these results, magnetostrictive stimulation was selected as the preferred approach for all subsequent *in vivo* experiments. This decision was further supported by the absence of cytotoxic effects in human brain tissue under the same stimulation conditions, validating the feasibility and safety of this strategy for functional neuromodulation.

### 2.3 Wireless Modulation of Motor Behavior in Freely Moving Mice

Based on *in vitro* results demonstrating that magnetostrictive stimulation elicits significantly stronger neuronal activation compared to torque-based actuation (Section 2.2), we selected the magnetostrictive regime for all *in vivo* experiments. This mode combines a static and 150 Hz alternating magnetic field to induce axial strain in the MEFs, enabling remote neuromodulation without invasive wiring. To establish a behavioral baseline and serve as an internal control, mice were first exposed to a static magnetic field (DC 120 mT) alone. One week later, the same animals were exposed to the full magnetostrictive stimulus (DC 120 mT + AC 6 mT at 150 Hz), allowing within-subject comparison of behavioral outcomes.

To evaluate functional efficacy, 1 mm^2^ sections of MEFs were surgically implanted into the left premotor cortex of freely moving mice (n = 8) (**Figure 5a**). The premotor cortex was selected based on previous findings demonstrating its capacity to elicit motor responses upon wireless stimulation[27]. Following a two-week recovery period, animals underwent behavioral testing under magnetic stimulation inside a solenoid coil.

**Figure 5.**
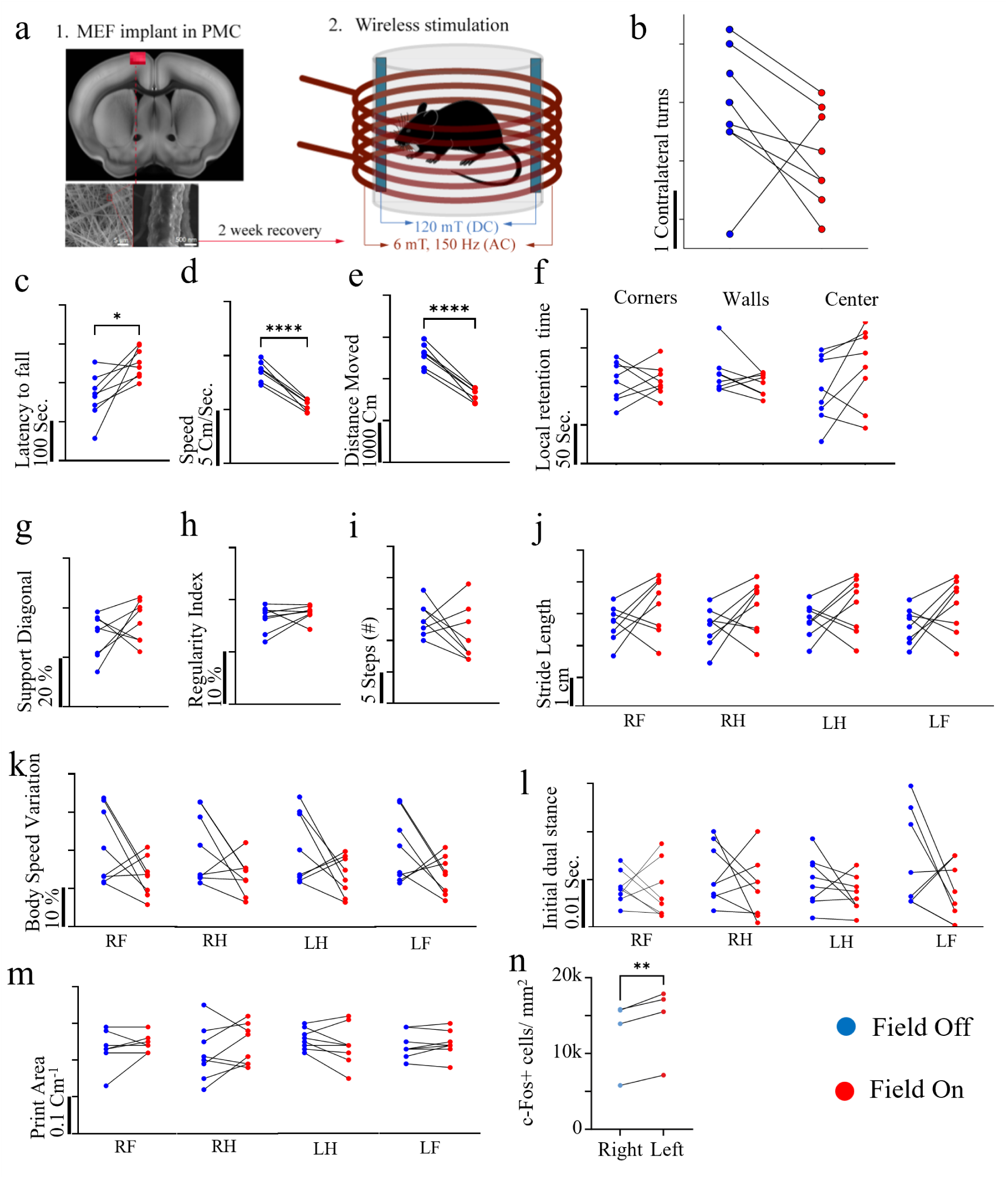
Magnetoelectric stimulation of the premotor cortex. a) Schematic of the implantation and stimulation setup. A 1 mm^2^ MEF patch is implanted into the left premotor cortex and activated using a combined static (DC 120 mT) and alternating (AC 6 mT, 150 Hz) magnetic field (n=8). c) Rotarod performance (latency to fall) under MEF-off and MEF-on conditions; mice show significantly enhanced motor coordination following stimulation (p < 0,05). d-e) Open field test results: total distance travelled, and average locomotion speed are significantly reduced after MEF stimulation, suggesting altered motor drive. Time spent in different arena zones remains unchanged, indicating no anxiety-related behaviour (p < 0.0001). g–m) CatWalk gait analysis reveals stimulation-induced trends: increased diagonal limb support (g), higher regularity index (h), reduced number of steps (i), increased stride length (j), decreased body speed variation (k), reduced initial dual stance time (l), and smaller paw print area (m). n) Increased c-Fos expression in the PMC implantation area corroborates the effect of the stimulation mediated by wireless MEF approach. (p<0.0018). RF: right front paw, RH: right hind paw, LF: left front paw, LH: left hind paw.

In the first test, mice were exposed to a static magnetic field only (DC 120 mT) for 3 min, and spontaneous rotations were recorded. The test was repeated one week later under combined static and alternating magnetic fields (DC 120 mT + AC 6 mT at 150 Hz). No significant difference in leftward rotations was observed between conditions (Figure 5b), indicating that this specific behavioral measure was not affected by magnetic stimulation.

To probe broader motor effects, animals were subjected to a behavioral test battery. In the rotarod assay, MEF-stimulated mice showed a significant increase in latency to fall, indicating improved balance and motor coordination (Figure 5c). In the open field test, magnetic stimulation led to a significant reduction in both locomotor speed and total distance traveled (Figures 5d–e), while spatial distribution across arena zones remained unchanged (Figure 5f), ruling out anxiety-driven effects.

Gait analysis using the CatWalk system revealed consistent trends across multiple coordination-related parameters. MEF stimulation was associated with increased diagonal limb support (Figure 5g), higher regularity index (Figure 5h), reduced step count (Figure 5i), longer stride length (Figure 5j), reduced body speed variation (Figure 5k), and shorter initial dual stance time (Figure 5l). A reduction in paw print area was also observed in several animals (Figure 5m), suggesting alterations in gait mechanics. While not all parameters reached statistical significance, the convergence of multiple trends such as altered stride length, reduced body speed variation, and increased regularity suggests a consistent neuromodulatory effect on locomotor output. Interestingly, these behavioral findings were accompanied by increased c-Fos expression in the ipsilateral premotor cortex of stimulated animals compared to the contralateral side (Figure 5n, Figure S9), indicating localized neuronal activation in response to MEF stimulation.

Similar behavioral shifts have been reported by Fornia et al[28], who observed unconscious motor arrest during direct electrical stimulation of the human premotor cortex. This supports the interpretation that PMC-targeted MEF activation can influence motor output even in the absence of conscious volitional control. Together, these findings reinforce the translational potential of magnetoelectric stimulation for wireless, minimally invasive neuromodulation targeting motor circuits.

## 3. Conclusion

We present a fully organic magnetoelectric (ME) platform composed of electrospun PVDF nanofibers embedded with anisotropic magnetite nanodiscs (MNDs), enabling wireless neural stimulation via magnetostrictive actuation. The MND geometry facilitates effective strain transfer, while electrospinning preserves the piezoelectric β-phase of PVDF, yielding a magnetoelectric voltage coefficient of 1.26 V·cm⁻¹·Oe⁻¹, comparable to or exceeding values reported for hybrid and ceramic-based ME systems. *In vitro* studies confirmed that only high-frequency magnetostrictive stimulation elicited robust neuronal activation, with responses suppressed by tetrodotoxin, indicating voltage-gated ion channel engagement.

Biocompatibility of MEFs was validated e*x vivo* in human brain tissue which under repeated stimulation showed no increase in cell death. Wireless stimulation of the premotor cortex in freely moving mice revealed enhanced motor coordination and consistent trends in gait modification. These findings align with clinical reports of motor arrest during direct electrical stimulation of the premotor cortex in humans[28], supporting the functional targeting achieved via MEF activation.

Despite these promising results in neuromodulation, several limitations should be acknowledged. The behavioral effects observed *in vivo*, while consistent, did not reach statistical significance across all parameters, likely due to limited sample size. In contrast to conventional deep brain stimulation, which requires fully implanted wired systems, both MEF implants and nanoscale neuromodulation systems[14, 17, 29, 30] offer the advantage of wireless operation. MEF implants, in particular, offer several benefits over both nano- and macro-scale systems, including suitability for cortical applications. While cortical stimulation techniques such as motor cortex stimulation are already in clinical use, MEF implants provide a less invasive and more flexible platform with improved structural integrity, mechanical flexibility and potential for further miniaturization. However, the current magnetic setup requires relatively high static field strengths, which may limit immediate clinical translation, so the material functional properties and field amplitudes should be optimized simultaneously. Though PVDF is for long know material used in clinical applications[31], long-term stability and chronic performance of the MEF implants in the brain remain to be investigated in details.

Ultimately, this work provides the first demonstration of soft, biocompatible ME fibers enabling wireless neuromodulation in behaving animals. These findings lay the groundwork for scalable, tether-free neural interfaces that avoid the drawbacks of rigid implantable electronics and offer new opportunities in minimally invasive bioelectronic medicine.

## 4. Experimental Section

### 4.1 MEFs synthesis

Magnetoelectric nanofibers are generated by dispersing 640 mg of PVDF in a mixture of 1.77 mL of DMF and 1.15 mL of acetone, with continuous magnetic stirring at 50°C overnight.

A second reaction vial is hermetically sealed with a rubber stopper, and the mixture is collected with a syringe. Subsequently, it is injected into a dispersion containing 1 mg of MNDs, produced following protocol[14], in 1 mL of acetone, all while undergoing continuous sonication at a temperature of 50°C. The resulting dispersion is then subjected to electrospinning using an electrospinner (Fluidnatek LE-10 by Bioinicia), employing a high voltage of 15 kV, with a flow rate set at 1000 μL/h. The collector is positioned 15 cm from the ejection needle, and the electrospinning process lasts for 1 hour.

#### 4.1.1 MEFs Characterization

The characterization of the fabricated nanofibers is performed using a combination of morphological, structural, and spectroscopic techniques to assess their physical and chemical properties.

SEM Analysis: Scanning electron microscopy (SEM) of the magnetoelectric fibers (MEFs) and control nanofibers is performed using a field-emission SEM (Jeol JSM-F100) equipped with energy-dispersive X-ray spectroscopy (EDX). The instrument operates in low-pressure mode and features STEM capability, along with a sample plasma cleaner, to ensure optimal surface imaging.

XRD Analysis: X-ray diffraction (XRD) measurements are conducted on fiber samples using a Bruker D8 Advance diffractometer equipped with a Lynxeye XE-T detector using Cu Kα radiation (1.54 Å) in the 20 to 80 2θ angle diffraction range.

FTIR Spectroscopy: Fourier-transform infrared (FTIR) spectroscopy is carried out using an Alpha II compact FT-IR spectrometer. Measurements are taken in ambient air at room temperature, and transmittance spectra are recorded in the range of 400 to 4000 cm⁻¹. A platinum attenuated total reflectance (ATR) module with a diamond crystal is used, and each sample is carefully positioned on the crystal surface for consistent analysis.

Magnetometry: Magnetization of the samples is characterized in a SQUID magnetometer MPMS3 (Quantum Design) in a vibrating sample magnetometry (VSM) mode at 300 K.

#### 4.1.2 Magnetoelectric Coefficient Determination

The magnetoelectric coefficient of the magnetoelectric nanofibers was computed via the lock-in method in the frequency domain. The perturbation is driven by an AC power source (GW Instek ASR) and subsequently recorded via a precision lock-in amplifier (MFLI, Zurich Instruments). Electrospinning of the fibres was carried out on adhesive films and co-planar copper electrodes were attached with conductive silver paste. An AC perturbation of 120 Hz was applied via a Helmholtz which exhibited a self-resonance frequency of 250 kHz with the strength of the magnetic field varying linearly with the applied voltage. The magnetoelectric voltage coefficient was determined using Equation 1, which is based on the slope of the output voltage, δV, as a function of the frequency-dependent AC magnetic field, δH(f), normalized by the average nanofiber thickness, tPE. Concentric Helmholtz coils were custom-built to generate the AC perturbation required to measure the magnetoelectric (ME) root mean square voltage output. The magnetic flux density produced by these coils is calculated using Equation (2), and a schematic representation is provided in Figure S5. In this equation, μ_0_ denotes the permeability of free space, N represents the number of coil turns, I is the coil current, and R_1_ is the coil radius. The scaling factor, c_1_, which depends on the ratio of the coil radius to the gap separating the two coils, has a value of 0.45 for this experiment.

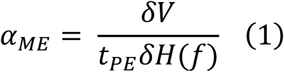

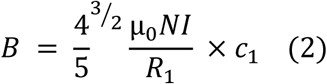

The baseline electromechanical noise of the system was accounted for by subtracting the output of the control PVDF nanofibers from the composite magnetoelectric fibres. To gauge the suitability of the method, the magnetoelectric coefficient was computed for variable AC magnetic fields at a fixed frequency and DC field. According to equation 1, the magnetoelectric voltage coefficient must not vary with AC field as the output voltage of the fibres varies linearly with the applied AC field leading to a fixed ratio pertaining to the coefficient. The tested fields consist of 3.2 Oe, 48 Oe and 80 Oe leading to an α_ME_ of 1.25 V/cm Oe, 1.20 V/cm Oe, and 1.3125 V/cm Oe confirming appropriate baseline noise removal.

### 4.2 Primary cortical neuron culture

Primary cortical neurons were isolated from P0–P2 Wistar rat pups under approved protocols (FAU Animal Welfare Committee, license TS 10-2023). Following dissection, cortical tissue was enzymatically digested in papain and gently triturated to yield a single-cell suspension. Cells were resuspended in Neurobasal medium supplemented with B-27 and Glutamax and seeded at 275,000 cells per well in 12-well plates containing coverslips pre-coated with either poly-D-lysine (PDL), standard nanofibers (NFs), or magnetoelectric nanofibers (MEFs). Cultures were maintained for 10 days, with one-third medium exchange every 3 days. To limit glial proliferation, 5-fluoro-2’-deoxyuridine (FUDR) was added between days 3 and 6.

### 4.3 *In vitro* calcium imaging

At day 10, cortical cultures (PDL (control), NFs, and MEFs) were incubated with 5 µM Fluo-4 and 10 µM Dooku1 inhibitor for 30 min to avoid potential neuronal mechanosensitivity arising from PIEZO1. Coverslips were additionally mounted upside-down in a Tyrode’s buffer - filled chamber using a 3D-printed spacer to minimize mechanical artifacts. Magnetic setups for stimulation are shown in photographs in Figure S7. For testing magnetoelectric regime, magnetic stimulation was delivered via a large-bore solenoid (5 kg copper spool, SchneiTec) powered by a Crown DC-300 amplifier receiving a 150 Hz, 5 Vpp signal and providing field amplitude of 5 mT. A 220 mT static field was generated using two neodymium N52 magnets (MagnetMax) positioned 3 cm apart inside the solenoid, with a 3D-printed holder, and fixed in place with epoxy resin. For testing torque regime, magnetic stimulation was delivered from similar coil, but without permanent magnets and with input signal 3.3 VP-P at 5 Hz, resulting in the generation of a magnetic field amplitude of 26 mT. Magnetic field values were verified using an SMP3 EMF meter (Wavecontrol). Four conditions were tested (n = 9 per group): PDL control, neurons on NFs, neurons on MEFs, and neurons on MEFs treated with 1 µM tetrodotoxin (TTX) to block voltage-gated sodium channels. For all recordings, three 20 s stimulation periods were applied within a 130 s imaging window. Videos of calcium activity were collected on an inverted Olympus IX73 microscope in the .vsi format using the Olympus cellSens software at the frame rate 4 Hz.

#### 4.3.1 Calcium Imaging Analysis

The analysis of calcium activity was based on a processing by Gregurec et al.[14] The image sequences were processed in ImageJ (BioFormats plugin). Regions of interest (ROIs) were segmented using thresholding and watershed algorithms, and mean fluorescence intensities were exported to MATLAB. ΔF/F_0_ traces were computed using a sliding window baseline correction (τ_1_ = 5, τ_2_ = 10 frames) and smoothed (τ_0_ = 3 frames) using an exponential moving average. Trace lengths were standardized, and 15 cells were randomly selected per trial. Activity thresholds were defined as 10× the baseline noise, and activity plots were generated as raster maps and population traces. Responses were further evaluated against a 5× baseline threshold to quantify active cell percentages over time. Significant population-level activity is further assessed by comparing each cell’s trace to a threshold of 5× the baseline standard deviation. The percentage of active cells per time point is calculated, plotted, and saved to Excel along with raw counts.

### 4.4. Biocompatibility in human brain slices and HEK293 cell line

The biocompatibility of material is tested on both HEKK cells 293 (Figure S8), and *ex vivo* on sections of human brain tissue obtained from epileptic patients and according to the protocol from Schwarz et al[32]. On the day of surgical resection, the tissue is maintained viable and sectioned into 300 µm slices using a vibratome Leica VT1200, Leica Biosystems, Wetzlar, Germany. These slices are then divided into three experimental conditions: (1) control, (2) NFs, and (3) MEFs. Each condition is subsequently subjected to stimulation by applying a direct current (DC) magnetic field of 120 mT and an alternating current (AC) magnetic field at 150 Hz and amplitude of 6 mT. After 24-hour incubation in artificial cerebrospinal fluid (aCSF), propidium iodide (PI) staining is performed to assess the number of dead cells resulting from the treatment. The second propidium iodide treatment is conducted after an additional 24 hours. The samples are imaged on a Leica DIVI LFS microscope (Leica Biosystem, Wetzlar, Germany) with a 20x water-dipping objective and images are processed in ImageJ to quantify fluorescence signal. The fluorescence signal density is calculated as Integrated Density normalized to the region of interest (ROI) area (IntDen/mm^2^). Quantification is performed on multiple non-overlapping fields per slice and per condition and background correction is applied uniformly if nonspecific signal is detected.

### 4.5 In-vivo study

#### 4.4.1 Subjects and housing

Adult male C57BL/6 mice (10 weeks, Charles River) were housed under standard conditions with ad libitum access to food and water (12:12 h reversed light/dark cycle). All procedures were approved by the Ethics Committee for Animal Experimentation, Maastricht University, in compliance with EU Directive 2010/63/EU.

#### 4.5.2 Surgical Procedure

A 1 mm^2^ section of MEF was implanted into the left premotor cortex of all mice using a stereotaxic apparatus (Stoelting, Wood Dale, IL, USA; model 51653). Mice were anesthetized with isoflurane (IsoFlo®, Abbott Laboratories Ltd, Berkshire, UK) and positioned in the stereotaxic frame with continuous inhalation anesthesia (2% isoflurane in 400 ml/min oxygen and 600 ml/min nitrous oxide) delivered via a nose cone. After exposing the skull with a scalpel, a 1 × 1 mm craniotomy was performed over the left premotor cortex (coordinates: 0-1 mm anterior to bregma; 0-1 mm lateral to bregma). The MEF was carefully placed on the cortex, after which the craniotomy was sealed with a thin layer of bone wax (Fine Science Tools, Heidelberg, Germany), and the skin was sutured. Mice were allowed to recover for two weeks.

#### 4.5.3 Experimental design

After recovery, mice underwent behavioral testing with and without magnetic field exposure. To reduce learning effects, rotarod and open field test conditions were counterbalanced and separated by four weeks. Prior to testing, mice were habituated for one week by daily placement in the inactive magnetic coil. Behavioral assessments began with mice placed in a transparent cylinder inside a copper solenoid coil containing two permanent magnets, generating a 120 mT DC field (as in Figure S7a, c). Mice remained in the coil for three minutes while their turning behavior was recorded and compared between field-on and field-off conditions, anticipating an increase in leftward turns during stimulation.

#### 4.5.4 Open field (OFT)

The open field test (OFT) took place in a square arena with 25 cm high walls and dimensions of 40 cm by 40 cm. A behavior tracking system, equipped with specialized software (EthoVision XT 15, Noldus Information Technology, Wageningen, The Netherlands), monitored each mouse. The behavioral recording lasted for 5 minutes.

#### 4.5.5 Rotarod

A rotarod with an accelerating grooved rotating beam (3 cm in diameter) elevated 16 cm above a platform (model 47650, Ugo Basile, Italy) was used to assess the coordination of the mice. The latency of falling from the rotating rod was recorded. Each mouse underwent four test sessions, each lasting 300 seconds, with and without AMF stimulation before the start. The test begins at a speed of 4 rpm and accelerates to 40 rpm over 300 seconds. To minimize stress and fatigue, mice were given at least 2 minutes of rest between trials. Results reflect the mean latency to fall from the rotarod across all test trials.

#### 4.5.6 Cat Walk

Mice were evaluated using the CatWalk-automated gait analysis system (CatWalkXT 10.6, Noldus). The setup included a long glass plate with fluorescent light projected onto the walkway floor from one side, allowing a camera mounted beneath the glass to capture the mouse’s footprints as it walks. Before each test, the glass plate was cleaned and dried to minimize the transmission of olfactory cues. A successful test recording typically consisted of an average of four uninterrupted runs with a consistent running speed and a maximum variation of 60%. The analysis included various parameters related to general movement, coordination, and static and dynamic gait patterns, such as run duration, number of steps, step pattern regularity index (%), stride length, number of step patterns, maximum contact area, print area, print length, print width, initial and terminal dual stance, body speed variation, three-limb support, and diagonal limb support.

### 4.6 Tissue collection and immunohistochemical analysis

At the end of the behavioral experiments, mice were divided into two groups: for *n* = 4 mice, the stimulation coil was activated, while for *n* = 4 mice, it remained off. Stimulation was applied for a total of 3 minutes, after which the animals were returned to their home cages. One hour later, mice were deeply anesthetized with an overdose of pentobarbital. Transcardial perfusions with PBS and then 4% paraformaldehyde were carried out. Brains were removed and placed in fresh fixative overnight at 4°C. Subsequently brains were transferred to 1% NaN3 at 4°C for long-term storage. For vibratome sectioning (Leica®, Wetzlar, Germany), brains were embedded in 10% gelatin from porcine skin (Sigma-Aldrich, Zwijndrecht, The Netherlands), and then cut into 30 µm slices in the frontal plane on a vibratome (Leica®, Wetzlar, Germany). Slices were immediately transferred into 1% NaN3 and kept at 4°C. For immunohistochemistry, sections were incubated overnight with polyclonal rabbit anti-c-Fos primary antibody at concentration of 1:1000 and followed by Alexa Fluor 568 goat anti-rabbit IgG (H+L)-polyclonal secondary antibody at a concentration of 1:200. The number of c-Fos positive cells was counted using the stereological procedure, optical fractionator. Counts were done using a microscope (Olympus® BX51W1), a motorized stage, and the StereoInvestigator software (MicroBrightField, Williston, VT). For c-Fos quantification, all positively labeled cells within the premotor cortex were counted in an average of three coronal sections, spaced 300 µm apart, using a 40× objective. The total number of positive cells was estimated based on the number of counted cells and the sampling probability and corrected for the analyzed area.

### 4.7 Statistical analysis

All data are represented as mean ± standard error of the mean (SEM), and analyses were performed with Graph Pad Prism 10.5.0 (774). Normality and homogeneity of variance of the data were checked using the Shapiro-Wilk test and normality plots. The in-vitro data was analyzed using one-way ANOVA for significance and Dixońs Q test for outlier detection using Origin 2024. Behavioral data were analyzed using paired-samples *t*-tests to compare magnetic field on versus off conditions within subjects. Stereological cell counts were analyzed using paired-samples *t*-tests comparing the left and right hemispheres within each subject and group. P-values <0.05 were considered significant.

### 4.8 Materials

Iron (III) Chloride Hexahydrate, Papain Neurobasal Media A+, B-27 supplement A+, Glutamax, Fluo4 and Alexa Fluor 568 goat anti-rabbit IgG (H+L)-polyclonal secondary antibody are purchased from Fisher Scientific, Sodium Acetate is purchased from Sigma Aldrich, PVDF with an average molecular weight (Mw ~534,000 GPC, powder) is purchased from Merck. DMF and acetone are sourced from Carl Roth, Polyclonal rabbit anti-c-Fos primary antibody is purchased from Abcam, and Tyrodés solution is prepared in-house following the protocol available in the supporting Information.

## Acknowledgments

L.S., V.D.T, E.K., N.I.G., F.W and D.G Acknowledge the European Innovation Council Pathfinder Challenge project, CROSSBRAIN (GA 101070908). D.G. and F.W. acknowledge ERC starting Grant BRAINMASTER (GA 101116410). We gratefully acknowledge the support of Prof. Dr. Karsten Meyer (Friedrich-Alexander-Universität Erlangen-Nürnberg; Lehrstuhl für Anorganische und Allgemeine Chemie) for providing SQUID magnetometer data.

## Data Availability Statement

The data that support the findings of this study are available from the corresponding author upon reasonable request.

## Author Contributions

L.S. and A.W. contributed equally to this work. D.G. SH, L.S., and A.W. conceived the idea. L.S. optimised the fabrication process, fabricated the samples and analysed physico chemical properties of the samples. J.E. and J.B contributed to the optimization of the fabrication process. L.S. and A.W. performed biocompatibility tests, *in vitro,* and behavioural analysis. V.D.T, F.W. and E.K. contributed to the data curation and analysis of structural properties. N.G. contributed to statistical analysis. M.S.B. performed ME characterisation and analysis. L.S., A.W., S.H. and D.G. drafted the manuscript, and all authors discussed the results and revised the manuscript.

## References

1. Liang, X., H. Chen, and N.X. Sun, Magnetoelectric materials and devices. APL Materials, 2021. 9(4).

2. Baji, A., et al., Electrospun barium titanate/cobalt ferrite composite fibers with improved magnetoelectric performance. RSC Advances, 2014. 4(98): p. 55217–55223.

3. Zheng, T., et al., Local probing of magnetoelectric properties of PVDF/Fe3O4 electrospun nanofibers by piezoresponse force microscopy. Nanotechnology, 2017. 28(6): p. 065707.

4. Tower, S.S., Arthroprosthetic Cobaltism: Neurological and Cardiac Manifestations in Two Patients with Metal-on-Metal Arthroplasty: A Case Report. JBJS, 2010. 92(17): p. 2847–2851.

5. Ikeda, T., et al., Polyneuropathy caused by cobalt–chromium metallosis after total hip replacement. Muscle & Nerve, 2010. 42(1): p. 140–143.

6. Rizzetti, M.C., et al., Loss of sight and sound. Could it be the hip? The Lancet, 2009. 373(9668): p. 1052.

7. Oldenburg, M., R. Wegner, and X. Baur, Severe Cobalt Intoxication Due to Prosthesis Wear in Repeated Total Hip Arthroplasty. The Journal of Arthroplasty, 2009. 24(5): p. 825.e15–825.e20.

8. Paralı, L., et al., Piezoelectric and magnetoelectric properties of PVDF/NiFe2O4 based electrospun nanofibers for flexible piezoelectric nanogenerators. Current Applied Physics, 2022. 36: p. 143–159.

9. Slavin, K.V., H. Nersesyan, and C. Wess, Peripheral Neurostimulation for Treatment of Intractable Occipital Neuralgia. Neurosurgery, 2006. 58(1): p. 112–119.

10. Lim, J.Y., S. Kim, and Y. Seo, Enhancement of β-phase in PVDF by electrospinning. AIP Conference Proceedings, 2015. 1664(1).

11. Ristić, D., P. Spangenberg, and J. Ellrich, Analgesic and antinociceptive effects of peripheral nerve neurostimulation in an advanced human experimental model. European Journal of Pain, 2008. 12(4): p. 480–490.

12. Bütefisch, C.M., et al., Enhancing Encoding of a Motor Memory in the Primary Motor Cortex By Cortical Stimulation. Journal of Neurophysiology, 2004. 91(5): p. 2110–2116.

13. Martins, P., A.C. Lopes, and S. Lanceros-Mendez, Electroactive phases of poly(vinylidene fluoride): Determination, processing and applications. Progress in Polymer Science, 2014. 39(4): p. 683–706.

14. Gregurec, D., et al., Magnetic Vortex Nanodiscs Enable Remote Magnetomechanical Neural Stimulation. ACS Nano, 2020. 14(7): p. 8036–8045.

15. Mascolo, M.C., Y. Pei, and T.A. Ring, Room Temperature Co-Precipitation Synthesis of Magnetite Nanoparticles in a Large pH Window with Different Bases. Materials, 2013. 6(12): p. 5549–5567.

16. Kim, S., et al., Utilization of a magnetic field-driven microscopic motion for piezoelectric energy harvesting. Nanoscale, 2019. 11(43): p. 20527–20533.

17. Kim, Y.J., et al., Magnetoelectric nanodiscs enable wireless transgene-free neuromodulation. Nature Nanotechnology, 2025. 20(1): p. 121–131.

18. Cai, X., et al., A critical analysis of the α, β and γ phases in poly(vinylidene fluoride) using FTIR. RSC Advances, 2017. 7(25): p. 15382–15389.

19. Jia, N., et al., Enhanced β-crystalline phase in poly(vinylidene fluoride) films by polydopamine-coated BaTiO3 nanoparticles. Materials Letters, 2015. 139: p. 212–215.

20. Li, L., et al., Studies on the transformation process of PVDF from α to β phase by stretching. RSC Advances, 2014. 4(8): p. 3938–3943.

21. Peng, Y. and P. Wu, A two dimensional infrared correlation spectroscopic study on the structure changes of PVDF during the melting process. Polymer, 2004. 45(15): p. 5295–5299.

22. Xin, Y., et al., Full-fiber piezoelectric sensor by straight PVDF/nanoclay nanofibers. Materials Letters, 2016. 164: p. 136–139.

23. Liang, C.-L., et al., Induced Formation of Dominating Polar Phases of Poly(vinylidene fluoride): Positive Ion–CF2 Dipole or Negative Ion–CH2 Dipole Interaction. The Journal of Physical Chemistry B, 2014. 118(30): p. 9104–9111.

24. Zhang, P., et al., Electrospun Doping of Carbon Nanotubes and Platinum Nanoparticles into the β-Phase Polyvinylidene Difluoride Nanofibrous Membrane for Biosensor and Catalysis Applications. ACS Applied Materials & Interfaces, 2014. 6(10): p. 7563–7571.

25. Sang, M., et al., Fabrication of a piezoelectric polyvinylidene fluoride/carbonyl iron (PVDF/CI) magnetic composite film towards the magnetic field and deformation bi-sensor. Composites Science and Technology, 2018. 165: p. 31–38.

26. Koç, M., et al., Piezoelectric and magnetoelectric evaluations on PVDF/CoFe2O4 based flexible nanogenerators for energy harvesting applications. Journal of Materials Science: Materials in Electronics, 2022. 33(10): p. 8048–8064.

27. Montgomery, K.L., et al., Wirelessly powered, fully internal optogenetics for brain, spinal and peripheral circuits in mice. Nature Methods, 2015. 12(10): p. 969–974.

28. Fornia, L., et al., Direct electrical stimulation of the premotor cortex shuts down awareness of voluntary actions. Nature Communications, 2020. 11(1): p. 705.

29. Signorelli, L., et al., Magnetic nanomaterials for wireless thermal and mechanical neuromodulation. iScience, 2022. 25(11): p. 105401.

30. Hescham, S.-A., et al., Magnetothermal nanoparticle technology alleviates parkinsonian-like symptoms in mice. Nature Communications, 2021. 12(1): p. 5569.

31. Laroche, G., et al., Polyvinylidene fluoride (PVDF) as a biomaterial: From polymeric raw material to monofilament vascular suture. Journal of Biomedical Materials Research, 1995. 29(12): p. 1525–1536.

32. Schwarz, N., et al., Human Cerebrospinal fluid promotes long-term neuronal viability and network function in human neocortical organotypic brain slice cultures. Scientific Reports, 2017. 7(1): p. 12249.

