## Supplementary information for "Biocompatible PVDF Nanofibers with Embedded Magnetite Nanodiscs Enable Wireless Magnetoelectric Neuromodulation"

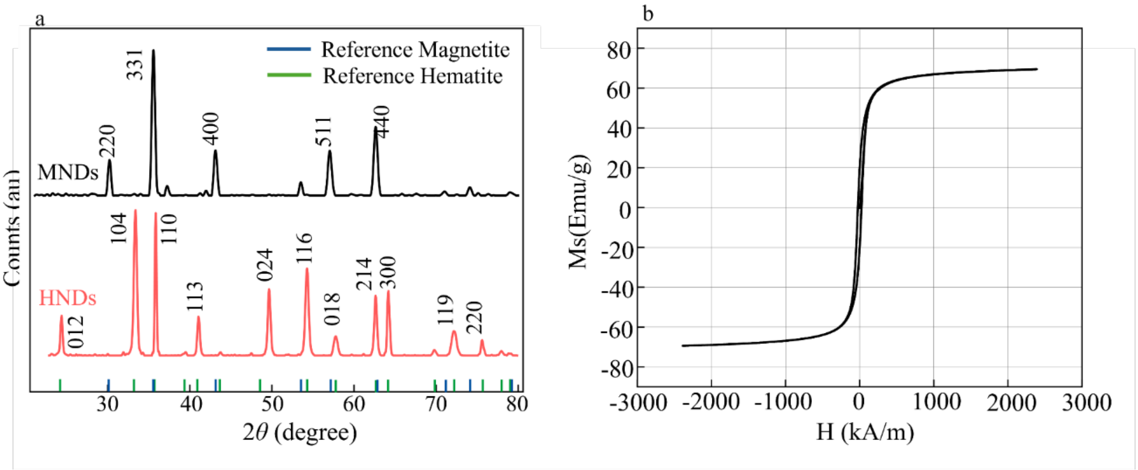


**Figure S1:** Magnetic nanodiscs material characterization a) XRD diffractograms of hematite nanodiscs (HNDs) and mangetite nanodiscs (MNDs) with references included, b) Saturation magnetization of MNDs (expressed as emu per gram of MND).

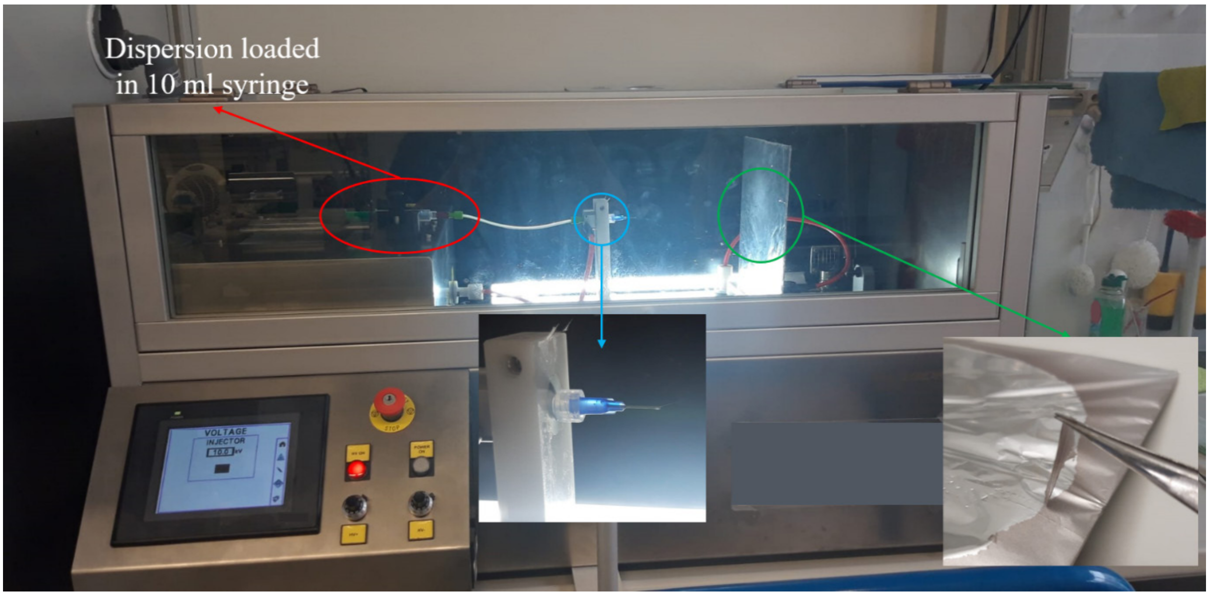


**Figure S2:** The dispersion of PVDF and nanodiscs is loaded into a 10 mL syringe and pushed through to a needle where a high voltage is applied. The extruded dispersion is then drawn toward the collector on the opposite side and stretched into a fiber. The resulting fibers are subsequently collected using tweezers.

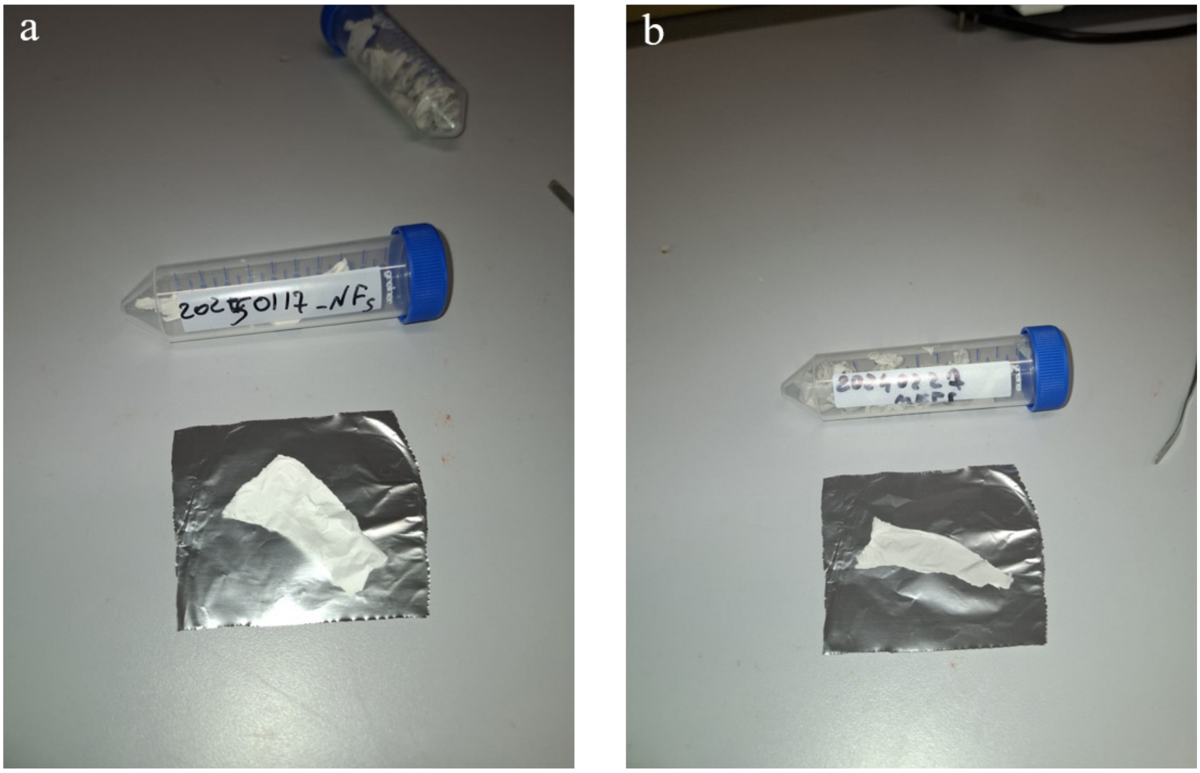


**Figure S3:** The fibers produced by electrospinning resemble a fabric; they are therefore very flexible and easy to handle.

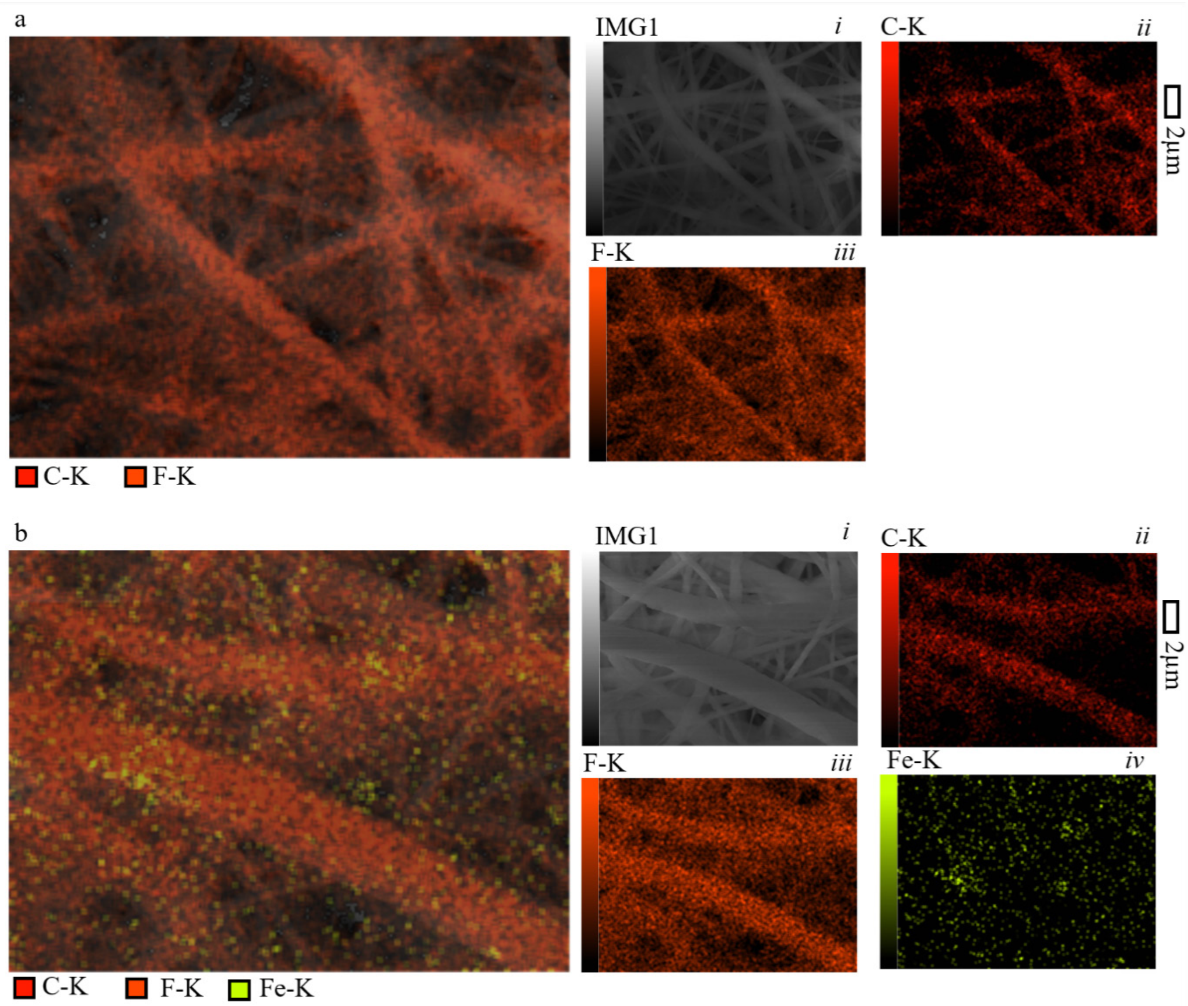


**Figure S4:** Elementary analysis by Energy Dispersive X-ray analysis (EDX) of nanofibres shows that: a) in pure PVDF nanofibres (NFs), combined with the corresponding SEM image (*i*), the two constituent elements of PVDF, carbon (*ii*) and fluorine (*iii*) are identified; b) in PVDF fibers enriched with MNDs, alongside the SEM image (*i*), in addition to carbon (*ii*) and fluorine (*iii*), the presence of iron (*iv*) is also detected, confirming the successful incorporation of MNDs.

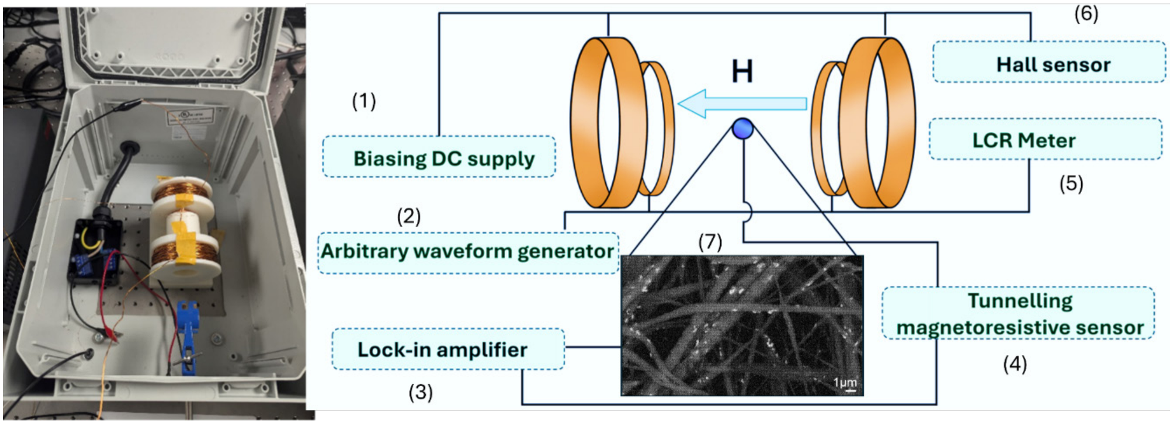


**Figure S5:** Setup and schematic diagram of the experimental setup used to characterize the magnetoelectric (ME) response of nanofibers. The setup includes: 1. A DC power supply or permanent magnets that provide a DC magnetic field in the plane of the nanofibers; 2. AC power generator that applies an AC field along the same plane; 3. Lock-in amplifier (Zurich Instruments) records the output voltage variation due to the ME fibers at the set frequency; 4. TMR sensor (Neuranics Ltd) for monitoring the AC magnetic field; 5. LCR meter (Keysight E4980AL) for monitoring the impedance of the AC coil to ensure that the coil functionality is not degraded at higher amplitudes; 6. Hall sensor (TLV493D) to measure the DC magnetic field; 7. SEM micrograph of MEFs

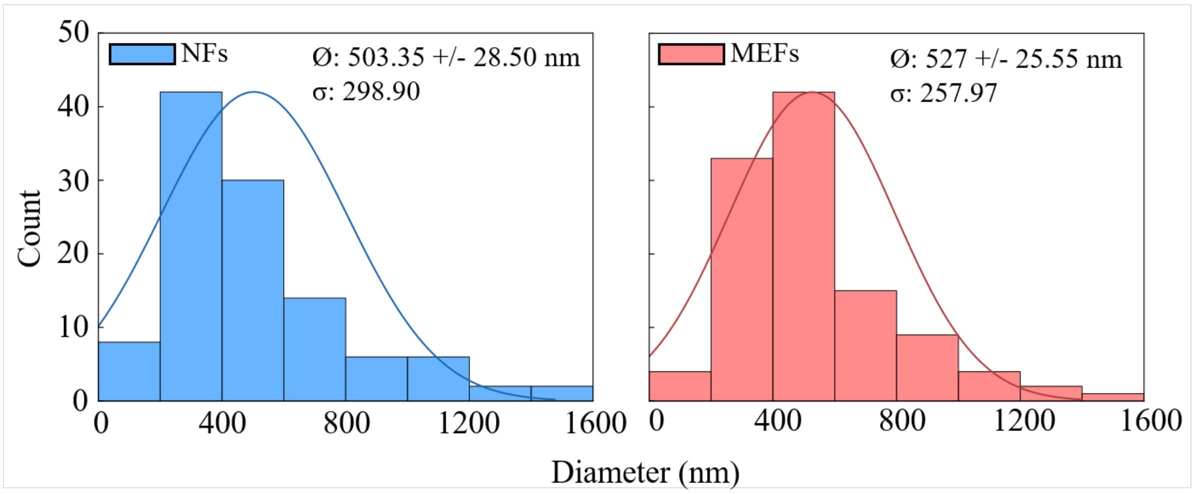


**Figure S6:** Distribution of the NFs and MEFs diameter.

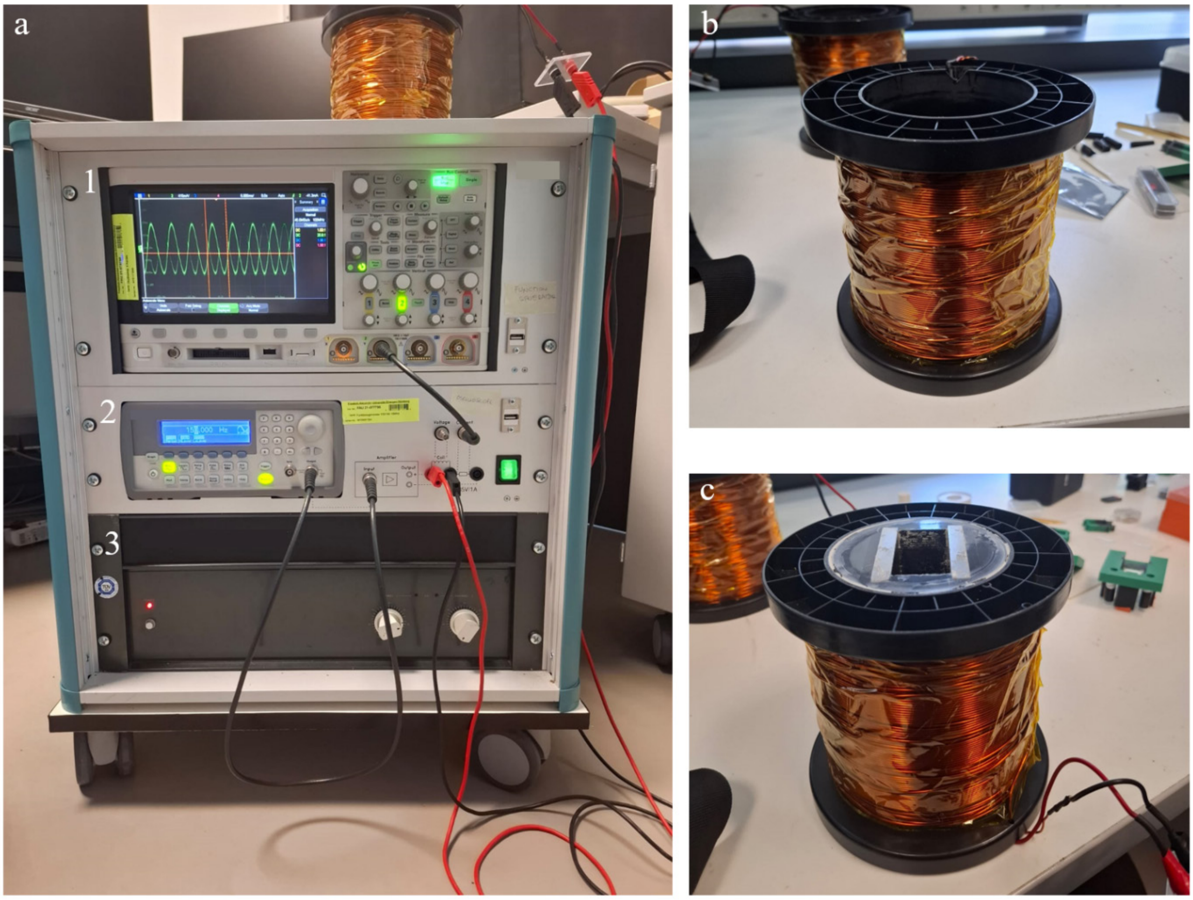


**Figure S7:** The setup used for stimulating the fibers during calcium imaging consists of: a) a main unit containing an oscilloscope (1), a wave generator (2), and an amplifier (3); this system is paired with b) a 5-kg copper coil for the magnetomechanical stimulation of the fibers; and for magnetoelectric stimulation, c) a 5-kg copper coil with two fixed neodymium permanent magnets inside, generating a constant magnetic field of 220 mT.

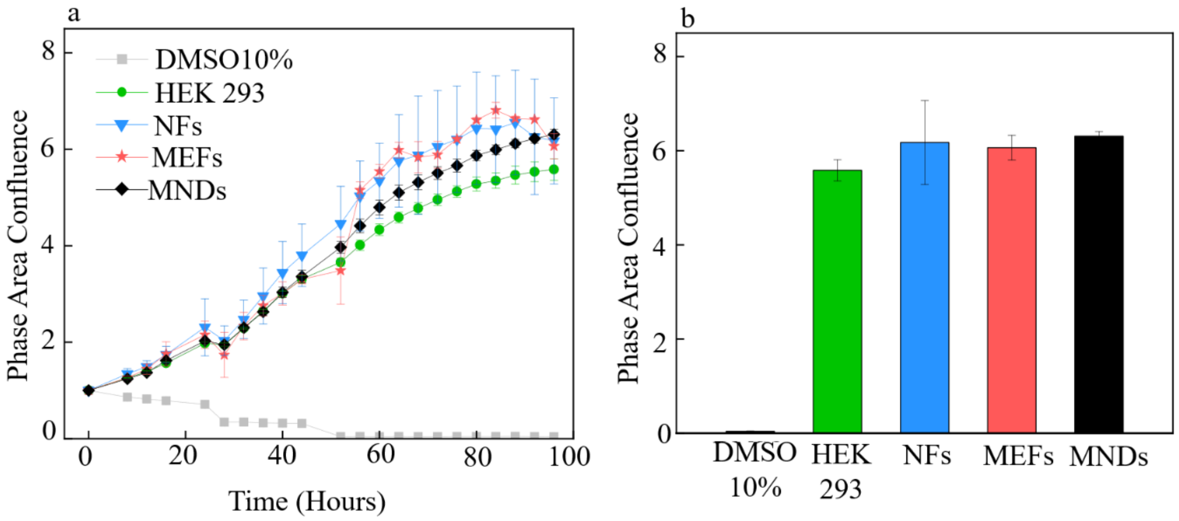


**Figure S8:** Phase area confluence of HEK293 cells: a) Variation of phase area confluence over a period of 100 hours in the presence of DMSO as a negative control, HEK293 cells as baseline control, NFs, MEFs, and MNDs; b) Phase area confluence at the end of the 100-hour period

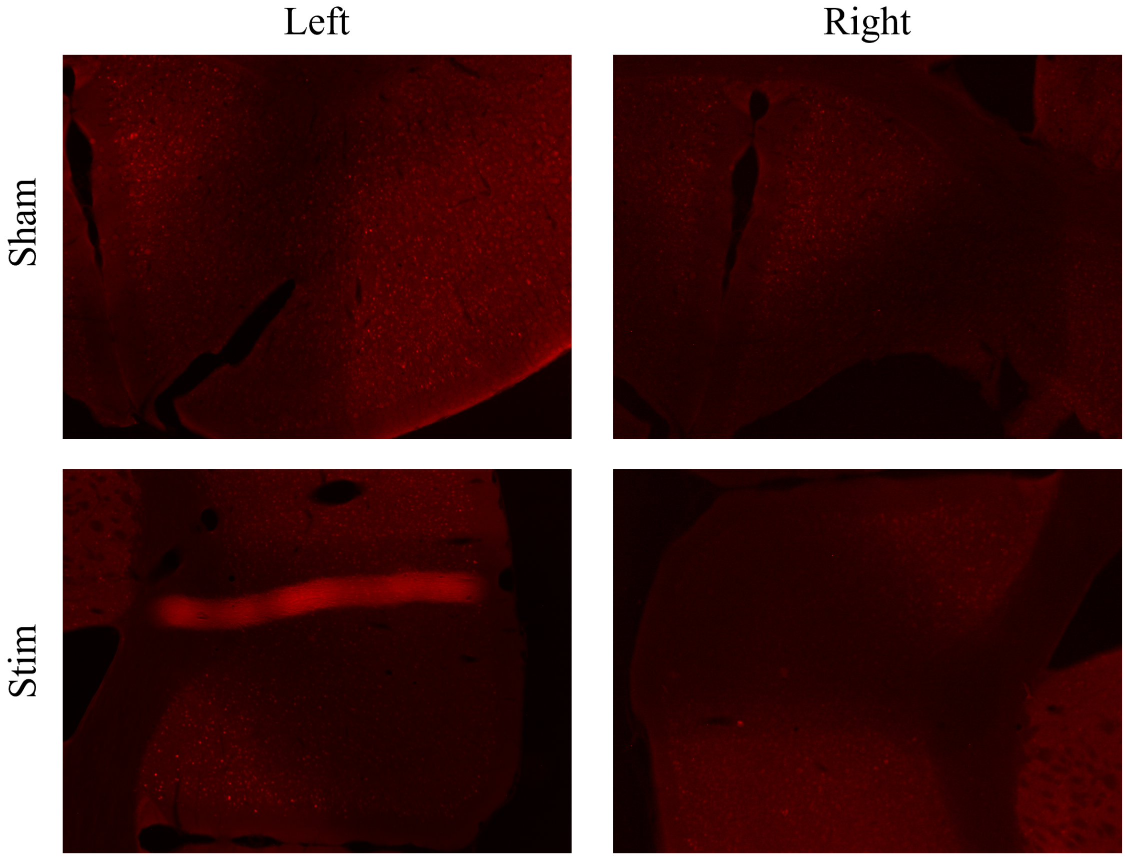

**Figure S9:** c-Fos staining (red) of left and right PMC in Sham (non stimulated, control) and in Stim (stimulated) WT mice demonstrating high expression of active neurons in the stimulated left PMC. Magnification 4x

**Table S1.** Tyrode´s Solution, HEPES-buffered

| Component | Final Working Concentration (1X) | Stock Concentration (10X) | Amount for 1L 10X stock |
|---|---|---|---|
| NaCl | 135 mM | 1.35 M | 78.95 g |
| KCl | 5 mM | 50 mM | 3.73 g |
| NaH2PO4 | 0.4 mM | 4 mM | 0.55 g |
| HEPES | 10 mM | 100 mM | 23.83 g |
| Glucose | 5.5 mM | 55 mM | 9.91 g |
| NaHCO3 (optional) | 12mM | 120 mM | 10.08 g |
| CaCl2 | 1.8 mM | (not included in Stock) | (add freshly) |
| MgCl2 | 1.0 mM | (not included in stock) | (add freshly) |

**Preparation of 10X Stock**

- Dissolve solid reagents (except CaCl2 and MgCl2) in ~800 mL deionized water while stirring
- Adjust pH to 7.4 using NaOH (1M) or HCl (1M)
- Bring volume to 1 L with deionized water
- Sterilize: filter through a 0.22µm filter (avoid autoclaving, as $Ca^{2+}$ may precipitate)
- Store at 4°C for up to a month or aliquot and freeze at -20°C for longterm storage

**Dilution for 1X Working Solution**

Before use, dilute 1:10 with deionized water and add freshly:

- CaCl to 1.8mM (1.8 mL of 1M CaCl2 per Liter)
- MgCl2 to 1.0 mM (1mL of 1M MgCl2 per Liter)
-

(This prevents precipitation of calcium salts in storage.)
